# Sensorimotor system engagement during ASL sign perception: an EEG study in deaf signers and hearing non-signers

**DOI:** 10.1101/558833

## Abstract

When a person observes someone else performing an action, the observer’s sensorimotor cortex activates as if the observer is the one performing the action, a phenomenon known as action simulation. While this process has been well-established for basic (e.g. grasping) and complex (e.g. dancing) actions, it remains unknown if the framework of action simulation is applicable to visual languages such as American Sign Language (ASL). We conducted an EEG experiment with deaf signers and hearing non-signers to compare overall sensorimotor EEG between groups, and to test whether sensorimotor systems are differentially sensitive to signs that are produced with one hand (“1H”) or two hands (“2H”). We predicted greater alpha and beta event-related desynchronization (previously correlated with action simulation) during the perception of 2H ASL signs compared to 1H ASL signs, due to greater demands on sensorimotor processing systems required for producing two-handed actions. We recorded EEG from both groups as they observed videos of ASL signs, half 1H and half 2H. Event-related spectral perturbations (ERSPs) in the alpha and beta ranges were computed for the two conditions at central electrode sites overlying the sensorimotor cortex. Sensorimotor EEG responses in both Hearing and Deaf groups were sensitive to the observed gross motor characteristics of the observed signs. We show for the first time that despite hearing non-signers showing overall more sensorimotor cortex involvement during sign observation, mirroring-related processes are in fact involved when deaf signers observe signs.

## 1. The human action observation system

In recent years there has been increasing interest in how perception can be modulated by past physical experience. This work primarily investigates patterns of brain activation during observation of different actions with the overarching objective of explaining how action experience affects subsequent action perception. Current cognitive neuroscience research suggests that the action observation network (AON) facilitates action prediction and understanding (Rizzolatti & Fogassi, 2014; Rizzolatti & Sinigaglia, 2010), contains ‘mirror neurons’ (i.e. Mirror Neuron System, MNS; Bowman et al., 2017; Debnath, Salo, Buzzell, Yoo, & Fox, 2019), and plays a large role in the development in human social cognition (Iacoboni, 2009; Rizzolatti & Sinigaglia, 2010). The AON encompasses regions in the premotor cortex, inferior parietal lobule, supplementary motor area, and primary somatosensory cortex (Caspers, Zilles, Laird, & Eickhoff, 2010; Cross, Hamilton, Kraemer, Kelley, & Grafton, 2009). Prior research suggests that physical experience with actions affects the functioning of these systems, but the exact nature of this effect is not fully understood.

In an effort to clarify whether AON activation is related to a person’s individual motor repertoire (Calvo-Merino, Glaser, Grèzes, Passingham, & Haggard, 2005), recent research is exploring the differences in AON involvement in trained experts versus untrained novices, or how experience modulates AON involvement. Studies are exploring the effects of experience on simple actions (i.e. grasping; Braadbaart, Williams, & Waiter, 2013; Drew, Quandt, & Marshall, 2015; Quandt & Marshall, 2014) and complex actions (e.g. dancing; Calvo-Merino et al., 2005; Cross, Hamilton, & Grafton, 2006; Gardner, Goulden, & Cross, 2015; Orgs, Dombrowski, Heil, & Jansen-Osmann, 2008). The AON has been the focus of a wide variety of research that exemplifies its far-reaching involvement in human action processing, such as: biological motion represented as point light displays (Ulloa & Pineda, 2007), gestures (Montgomery, Isenberg, & Haxby, 2007; Quandt, Marshall, Shipley, Beilock, & Goldin-Meadow, 2012), facial expressions (van der Gaag, Minderaa, & Keysers, 2007), and brain computer interfaces (Pineda, Allison, & Vankov, 2000). While research within this field has grown within the past 20 years, there remains much to be understood about the complicated relationship between action experience and action perception.

### 1.1. Experience and action mirroring

Some researchers have demonstrated a linear relationship between increased motor experience and increased MNS activation (Calvo-Merino et al., 2005; Cross, Hamilton, & Grafton, 2006a; Orgs et al., 2008) while others posit a more complex, non-linear relationship that is based on a gradient of prior personal experience that influences neural efficiency (Babiloni et al., 2009; 2010; Gardner, Aglinskas, & Cross, 2017; Hobson & Bishop, 2017). The hypothesis for a linear relationship between activation in the AON and experience postulates that the more experience a person has with an action, the greater the mirror mechanism involvement during observation of said action. This phenomenon is said to enable automatic comprehension of the action, thus facilitating understanding (Rizzolatti & Fogassi, 2014). Given that motor imagery, action observation, and movement execution utilize similar premotor-parietal and somatosensory networks (Hardwick, Caspers, Eickhoff, & Swinnen, 2017), it is logical to consider a dependent relationship between action observation and execution. This positive linear relationship has been demonstrated in studies of complex actions with experts and novices (Calvo-Merino et al., 2005; Cross et al., 2012; Orgs et al., 2008).

However, a growing body of evidence strongly suggests that this relationship cannot be fully explained by the direct matching theories of action observation (Cross et al., 2011; Gardner et al., 2017; Gazzola, Rizzolatti, Wicker, & Keysers, 2007; Liew, 2013). This work shows greater sensorimotor activity when participants are unfamiliar with the observed action, compared to actions they are familiar with. This increased sensorimotor activity during observation of unfamiliar actions is hypothesized to be an effect of heavy feedforward processing—the brain attempting and failing to match the observed action, consequently manifesting as robust AON activation (Gardner et al., 2017). Within this non-linear framework, studies explaining increased recruitment/experience through direct matching theories are not accounting for the spectrum of human action experience. Given that there may not be a linear relationship between experience and activation, a non-linear relationship that relies on a familiarity continuum ranging from no experience, to some experience, to much experience, may provide a more complete explanation (Cross et al., 2012). Thus, under this non-linear framework, it would be possible to see increased sensorimotor activation for those with almost no experience and those with an expansive amount of experience and lessened recruitment for those with moderate familiarity (Cross et al., 2012; Gardner et al., 2017; Liew, 2013).

These experience-dependent AON investigations, in addition to many others, attempt to further explain the nature of the relationship between action experience and action perception. Contention with direct matching theories of action observation demonstrate the need to further explore the plasticity of the sensorimotor system as it relates to action experiences. As this body of AON research continues to grow, there are only a handful of studies attempting to connect action observation to a population that utilizes action as its main form of communication: sign language users.

### 1.2. American Sign Language: signs as actions

While it has been demonstrated that the AON is responsive to non-object oriented (intransitive) actions, pantomimes (Grèzes, Armony, Rowe, & Passingham, 2003), and communicative hand gestures (Montgomery et al., 2007), only recently have studies begun exploring to what extent sign language recruits the AON. In general, observation of sign language does not recruit the mirroring aspects of the AON for deaf fluent signers; a result that conflicts with the direct matching hypothesis (Fadiga, Fogassi, Pavesi, & Rizzolatti, 1995; Hari et al., 1998; Rizzolatti, Fogassi, & Gallese, 2001).

The involvement of the AON in sign language perception has been examined in a number of neuroimaging studies. In an functional neuroimaging (fMRI) study, Emmorey and colleagues (2010) found that Deaf signers do not activate the MNS while watching pantomimes (e.g. peeling a banana, running in place) or action verbs, while hearing signers recruited inferior frontal gyrus, premotor cortex, and the inferior/superior parietal lobule (areas associated with the MNS; Rizzolatti & Sinigaglia, 2010) under these same conditions. These results echo that of Corina et al. (2007) who found that Deaf signers did not significantly engage MNS areas during observation of different classes of human actions (i.e. self-oriented, object-oriented), while hearing non-signers reliably recruited frontoparietal areas during the same task. Other fMRI research indicates that signers recruit more traditional language and auditory areas and less MNS/AON areas during observation of either sign language or gesture compared to hearing signers or non-signers (MacSweeney et al., 2004; Newman, Supalla, Fernandez, Newport, & Bavelier, 2015). Interestingly, deaf non-signers have been found to activate the AON similarly during observation of communicative gestures and non-communicative gestures (Muise-Hennessey et al., 2016), suggesting that lack of MNS involvement for deaf signers is dependent on sign language knowledge, not auditory deprivation alone.

It has been theorized that non-signers activate the AON more strongly for gesture compared to signing, although both contain little to no linguistic information for this population. This pattern may be due to a hearing non-signer’s greater potential to imitate gestures, since they do not contain the same details, such as varying handshape and movement, as sign language does (MacSweeney et al., 2004). This notion is consistent with recent research in the match between an observer’s motor experience and the performer’s motor skill impacts the degree of AON involvement during perception (Errante & Fogassi, 2019). In contrast, deaf fluent signers are drawing upon both motor and linguistic knowledge when viewing signs. Signers may implicitly apply sign language knowledge when processing gesture or pantomime in an attempt to derive meaning from it, transferring the cognitive load to more traditionally linguistic areas (i.e. fronto-temporal areas).

While studies thus far have explored the AON in relation to sign language comparing: gestural communication systems (MacSweeney, Woll, & Campbell, 2002), pantomimes (Emmorey, Xu, Gannon, Goldin-Meadow, & Braun, 2010), and intransitive self-oriented actions vs. transitive object-oriented actions (Corina et al., 2007), no work has investigated sign language and the AON through a sign-only paradigm for both hearing and deaf participants. While previous work contrasts sign with gesture, sign language has yet to be further broken down into additional action categories, leaving it unknown if deaf signers recruit the AON differently for varying sign types. Historically, action observation research uses the same category of complex action for stimuli (i.e. dancing) but then breaks that down further (i.e. ballet or capoeira) to control for perceptible differences within the complex action itself (Calvo-Merino et al., 2005). Sign language research would benefit from being explored in the same manner as other action observation research, framing signs as complex actions and controlling for perceptible differences, as the AON may be differentially sensitive to the gross sensorimotor characteristics of observed signs.

### 1.3. Action Mirroring and EEG

The extent of desynchronization in the alpha (8-13 Hz) and beta (14-25 Hz) frequency ranges during action observation can indicate the degree of activity in sensorimotor aspects of the AON (Fox et al., 2016; Rizzolatti & Sinigaglia, 2010). Specifically, a frequency within the alpha range (8-13 Hz) over central electrode sites known as the “mu rhythm” has a particular sensitivity to sensorimotor stimuli and is a known indicator of MNS involvement (Arnstein, Cui, Keysers, Maurits, & Gazzola, 2011a; Bowman et al., 2017; Hari et al., 1998; Muthukumaraswamy, Johnson, & McNair, 2004).

When a person observes an action, sensorimotor cortices activate as if the observer is the one performing the action, a phenomenon referred to as direct matching. However, the action is not explicitly carried out by the observer, resulting in an inhibition of the action (i.e. action is only being simulated). This process results in desynchronization of alpha and beta rhythms over the sensorimotor cortex (Fox et al., 2016; Hari et al., 1998).

Previous work exploring the AON in relation to sign language use has largely used fMRI and PET techniques. While imaging modalities such as these are valuable in recognizing areas of the brain that are involved in higher level cognitive processes, including that of the AON, other techniques such as electroencephalography (EEG) can provide information unable to be gathered using other neuroimaging methods. Unlike fMRI or PET, EEG allows for very precise timing of activation related to observation of action (at the sacrifice of lessened spatial resolution; Fox et al., 2016). The mu rhythm occurs within alpha range frequencies at around 8 – 13 Hz (Bowman et al., 2017; Hari, 2006; Muthukumaraswamy et al., 2004; Quandt et al., 2012; Quandt & Marshall, 2014; Quandt, Marshall, Bouquet, & Shipley, 2013). A decrease in amplitude, or desynchronization, of these frequencies recorded over central and parietal electrode sites during action execution and observation reliably indexes sensorimotor cortical activation (Bowman et al., 2017; Fox et al., 2016). Recent work concluded that the EEG mu rhythm provides a valid index for the study of human neural mirroring (Debnath et al., 2019; Fox et al., 2016). Thus far, neuroimaging results in relation to sign language perception and the AON are not in line with other action observation research, as the action experts (i.e. signers) do not robustly recruit the MNS. By investigating this topic through the mu rhythm, our results will inform the relationship between sign language perception and MNS involvement while also adding to the growing body of evidence that investigates the overall relationship between experience and AON recruitment.

### 1.4. The current study

By framing sign language research as action, we aim to shed light on the role the AON plays in sign language perception for users and non-users of the language, resulting in a better understanding of overall AON functioning. We aimed to answer two questions: 1) How do Deaf and Hearing groups differ in sensorimotor EEG rhythm activity during perception of ASL signs; and 2) are both groups sensitive to the gross sensorimotor characteristics of observed signs? The second of these questions allows us to address the possibility that even if signers show less activity in AON regions during action observation, they may still be drawing upon their own sensorimotor systems when perceiving signs. To answer these questions, we designed an EEG study in which Deaf signers and Hearing non-signers both watched videos of ASL signs which varied in their sensorimotor demands: half of the videos showed signs that used one hand, while half of the signs used two hands.

## 2. Materials and Methods

### 2.1. Participants

Participants were recruited from Gallaudet University and surrounding Washington, D.C. communities through flyers, Craigslist, and Facebook advertisements. All deaf participants were self-identified as fluent in American Sign Language. All Hearing participants reported typical hearing and their sign language familiarity on a 1-10 scale. We excluded participants if they rated their own skills a 3 or above. Participants signed an informed consent form presented in written English and/or American Sign Language that had been approved by the University Institutional Review Board. Participants were compensated $20.00 an hour for their time.

Educational, demographic, and language background information is shown in Tables 1 and 2. Data from all participants in the current study was also analyzed in another recent publication from our laboratory (Quandt & Kubicek, 2018).

**Table 1.**
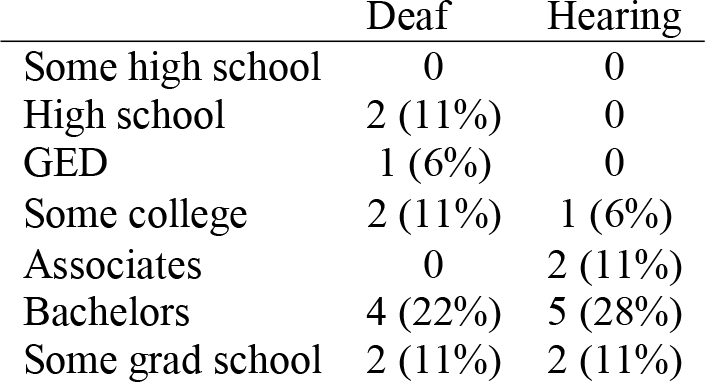
Formal education. Self-reported highest educational degree obtained for Deaf and Hearing participants. Percentages were rounded to the nearest whole number.

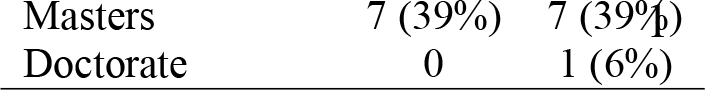

**Table 2.**
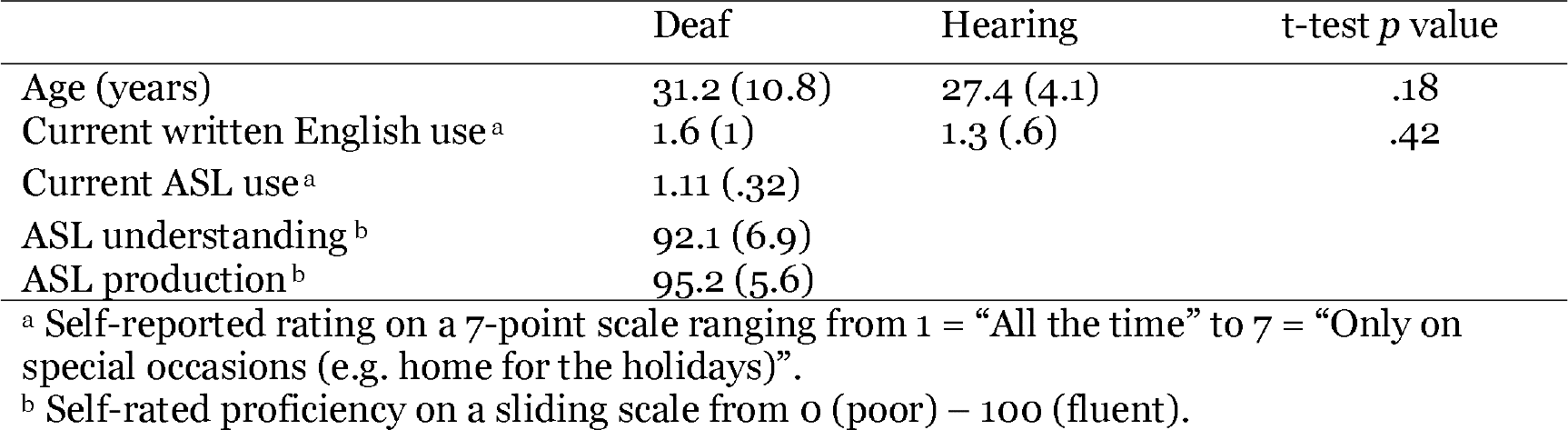
Demographics and language background information for all participants.

### 2.2. Stimuli

All stimulus videos were selected from ASL-LEX: A Lexical Database for ASL (www.asl-lex.org; (Caselli, Sehyr, Cohen-Goldberg, & Emmorey, 2016) and presented in E-Prime 2.0 (Psychology Software Tools). Two types of videos were used, with 40 of each of type. One video type consisted of signs that only use one hand (‘one-handed’, ‘1H’; i.e. GUILT). The other video type consisted of signs that use two hands (‘two-handed, ‘2H’; i.e. FAMILY). Of the 40 two-handed signs, 25 were symmetrical. There were no significant differences between the signed 1-handed (1H) or 2-handed (2H) words or their English translations for any of the following measures: frequency, iconicity, flexion, phonological properties, imageability, sign onset time (ms), sign offset time (ms) and sign length (ms; for more information see Caselli et al. 2017). Signs selected were nouns, adjectives, and adverbs (no verbs were included in the stimuli set). 1H and 2H signs did not significantly differ (*p* > .05) on overall total duration of signs or time of sign onset. Total clip length was significantly different (*p* < .05) for 1H (M = 1658.3 ms, SD = 415.5) and 2H (M = 1898.6 ms, SD = 480.6) signs. Videos consisted of a native signer producing each individually signed word, separated by presentation of a fixation point (duration = 1-2.5 s). Sign onset was the time of the first video frame where a fully-formed handshape contacted the body. If the sign did not include contact, sign onset was considered to be when the handshape arrived at the target location near the body or in neutral space (Caselli et al., 2016). Participants were told to keep track of how many “freeze frames” (stilled frames within the presented video) they saw within the presentation of 92 videos (12 freeze frames total).

**Table 3.** English and ASL norms for two categories of Stimuli. 1-handed (1H) words and 2-handed (2H) words, as in Quandt & Kubicek, 2018.

|  |  | **1H words: M**<br>**(SD)** | **2H words: M**<br>**(SD)** | **t-test *p***<br>**value** |
| --- | --- | --- | --- | --- |
| English norms | Word length | 5.2 (1.7) | 5.7 (2.2) | .25 |
|  | Log frequency (SUBTLEX) | 5.6 (10.6) | 14.3 (43.6) | .22 |
|  | Log frequency (HAL) | 9.6 (1.7) | 10.1 (1.5) | .24 |
|  | # phonemes | 4.1 (1.1) | 4.6 (1.3) | .10 |
|  | Lexical Decision RT M | 605.6 (45.5) | 616.2 (64.4) | .40 |
|  | Lexical Decision RT SD | 202.1 (64) | 215.4 (89) | .45 |
|  | Naming RT M | 605.5 (42.6) | 607.8 (42.5) | .81 |
|  | Naming RT SD | 128 (58.1) | 130.3 (53.1) | .85 |
|  | Imageability | 509.5 (112.5) | 543.6 (79.6) | .18 |
| ASL norms | Frequency (ASL-Lex) | 4.7 (.8) | 4.6 (1.1) | .42 |
|  | Iconicity | 3.7 (1.7) | 3.8 (1.4) | .90 |
|  | Flexion | 3.4 (2.2) | 3.6 (2.2) | .69 |

### 2.3. Experimental design

18 hearing non-signing participants and 18 deaf signing participants were included in this analysis. While sitting, participants observed a presentation of a signed word given by a deaf fluent signer (videos stimuli from *asl-lex.org*; Caselli et al., 2016) on a computer monitor approximately 2.5 feet away. After presentation of the stimulus, a fixation cross was presented for 1 - 2.5 s (randomized) in the center of the screen. Each video clip contained an audio trigger at the moment of sign onset (as defined by the ASL-LEX database), which was recorded by the EEG recording software. Participants were told to keep track of how many “freeze frames” they saw within the presentation of 92 videos. Recordings from these ‘catch-trials’ were not analyzed at any point in time. The full experiment consisted of four blocks with a total of 92 stimuli. The 92 stimuli were randomized then separated into four blocks of 23 trials each. Each video stimulus was seen only once, and breaks were given in between blocks.

### 2.4. Procedure and Recording

Following the consent procedure and an introduction to the experiment, participants were fit with the appropriate EEG cap and completed a language background form. Participants were instructed to attend to “freeze-frame” trials as described above and asked to report to the experimenter how many of these trials they had seen after each block of videos. A practice portion consisting of 10 trials took place to ensure the participant’s understanding of the task.

EEG was recorded from 64 active Ag/AgCl electrodes using an actiCAP setup (Brain Products GmbH, Germany), in combination with SuperVisc electrode gel. The EEG signals were amplified by the individual electrode amplifiers, and again by a 24-bit actiCHAMP amplifier (Brain Vision LLC, Morrisville, NC). Hardware filter settings included a high-pass filter (.53 Hz) and a low-pass filter (120 Hz).

### 2.5. Data preparation

All data processing was implemented using EEGLAB v. 14.1.1 (Delorme & Makeig, 2004). Data were referenced offline to the average of the two mastoid electrodes (TP9 and TP 10). Data were filtered offline using a .1 Hz highpass and a 100 Hz lowpass filter. Epochs were extracted from the continuous EEG, time locked to the sign onset of each stimulus of interest (1H and 2H signs). Data was epoched at −1.5 s to 2.5 s, with 0 being the moment the sign begins. The epoch from - 1.5 to −1 sec was used as a baseline measure, during which time a fixation cross was on the screen. All epoched datasets were compiled into one study folder and assigned groups (Deaf and Hearing) and conditions (1H and 2H).

### 2.6. Predictions

Based on existing EEG research comparing action experts with novices, we predict that deaf fluent signers will show greater alpha and beta desynchronization over central electrodes compared to the non-signing Hearing group during sign observation. Increased involvement of sensorimotor cortices for experts would demonstrate greater sensorimotor resonance during action observation, a phenomenon suggested to facilitate greater action understanding. Alternatively, it is possible that deaf signers will show less desynchronization in sensorimotor EEG rhythms than hearing non-signers, as has been suggested by relevant work in fMRI (Emmorey et al., 2010). Our second prediction is that there will be a greater desynchronization of power observed at central electrode sites during observation of two-handed signs as compared to one-handed signs for both Deaf and Hearing groups due to greater vicarious engagement of the sensorimotor cortex. Execution of two-handed signs recruits primary sensory and motor cortices more strongly than one-handed signs (Emmorey, Mehta, McCullough, & Grabowski, 2016) a pattern consistent with non-linguistic actions (Toyokura, Muro, Komiya, Obara, 2002). Thus, we expect similar activations for the observation of one and two-handed signs. Alternatively, one or both groups may not be sensitive to the gross sensorimotor characteristics of observe signs, in which case there would be no difference in the degree of sensorimotor EEG desynchronization.

### 2.7. Planned Analyses

Planned t-tests were driven by *a priori* predictions developed before data analysis (see above). Tests included analysis of EEG waveforms across the scalp in four frequency bands: low alpha (8-10 Hz), high alpha (11-13 Hz), low beta (14-17 Hz) and high beta (18-25 Hz). For each of these frequency bands we divided the epoch of interest into four time bins, which covered the majority of the duration of the sign: 0-250 ms, 250-500 ms, 500-750 ms, and 750-1000 ms. In an effort to avoid spurious findings, we only considered an effect noteworthy if it was significant at three or more adjacent electrodes. To control for multiple comparisons, we used a corrected *p* value of 0.16 (.05/3) as the threshold for significance. This analytical approach has been used in previously published work (Quandt & Kubicek, 2018).

To more closely examine the time and frequency characteristics of sensorimotor cortex during observation of signs, we conducted time-frequency analyses at 21 electrodes over the pre- and post-central gyrus, comprising a central region of interest (ROI). The ROI included electrodes FC5, FC3, FC1, FCz, FC2, FC4, FC6, C5, C3, C1, Cz, C2, C4, C6, CP5, CP3, CP1, CPz, CP2, CP4, and CP6. At each of these electrodes time-frequency plots (from 0-1000 ms in time and 8-25 Hz in frequency, which includes both alpha and beta ranges) were compared between 1H and 2H signs for both the Deaf and Hearing groups. Effects within these electrodes are of particular interest to our investigation, as alpha and beta rhythms detected at central electrodes are associated with activity in pre- and post-central gyri (primary motor and primary somatosensory cortices, respectively), which are key regions of the AON (Arnstein, Cui, Keysers, Maurits, & Gazzola, 2011b; Perry & Bentin, 2009; Ritter, Moosmann, & Villringer, 2009). For ROI analyses, a *p* value threshold of .05 was used, with an FDR correction applied to control for false positives (Benjamini, Drai, Elmer, & Kafkafi, 1995).

## 3. Results

### 3.1. Full Scalp Analysis

In the lower alpha band (8-10 Hz), we found no significant (*p* < .016) differences between Deaf and Hearing groups in the 0-250 ms time bin. A significant (*p* < .016) main effect in the latter 3 time bins encompassing 250-1000 ms post-sign onset was found in left lateralized and bilateral parieto-occipital regions. During this time period, the Hearing group showed greater desynchronization in alpha power in compared to the Deaf group.

In the upper alpha band (11-13 Hz), similar to lower alpha, there were no significant differences between Deaf and Hearing groups in the 0-250 ms time bin (*p* > .016). The Hearing group showed significantly greater alpha desynchronization while observing signs as compared to the Deaf group in the latter three time bins encompassing 250-1000 ms post sign onset (*p* < .016). This effect was found in left central electrodes (see Figure 1).

**Figure 1.**
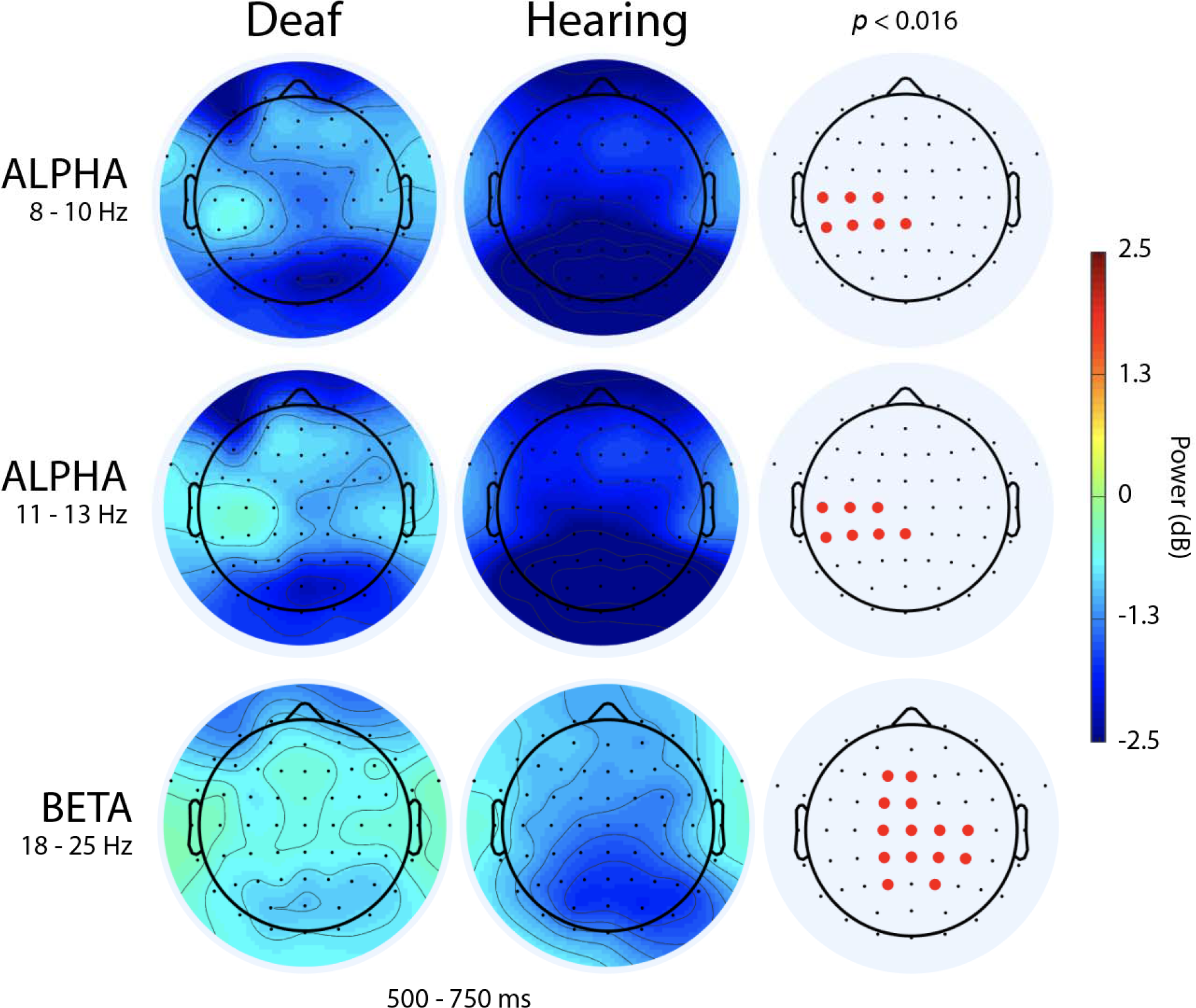
Full scalp maps of Deaf and Hearing groups in the 500 to 750 ms time bin. Lower (8-10 Hz) and upper (11 – 13 Hz) alpha range rhythms showing greater desynchronization for the hearing group in left-central electrodes. High beta range rhythm (18-25 Hz) showing greater desynchronization for the hearing group in bilateral central regions.

In the lower (14-17 Hz) and upper (18-25 Hz) beta band, all time bins (0-1000 ms) indicate significant (*p* < .016) differences between Deaf and Hearing groups in left frontal and central-parietal electrodes (see Figure 1) which indicates more beta desynchronization in the Hearing group when watching signs compared to the Deaf group.

### 3.2. Time-frequency analysis at ROI

In the Deaf group, six electrodes within the ROI showed significant effects of condition (1H v. 2H) on alpha power (8-13 Hz) during sign perception (*p* < .05, FDR corrected). Of these six significant effects, five indicated greater alpha desynchronization in the 2H condition as compared to the 1H condition (see Figure 2). In the remaining electrode, there was more alpha desynchronization for 1H. Eight electrodes showed significant effects of condition (1H v. 2H) on beta power (14-25 Hz) during sign perception (*p* < .05, FDR corrected). For all eight electrodes, there was greater beta desynchronization in the 1H condition as compared to the 2H condition.

**Figure 2.**
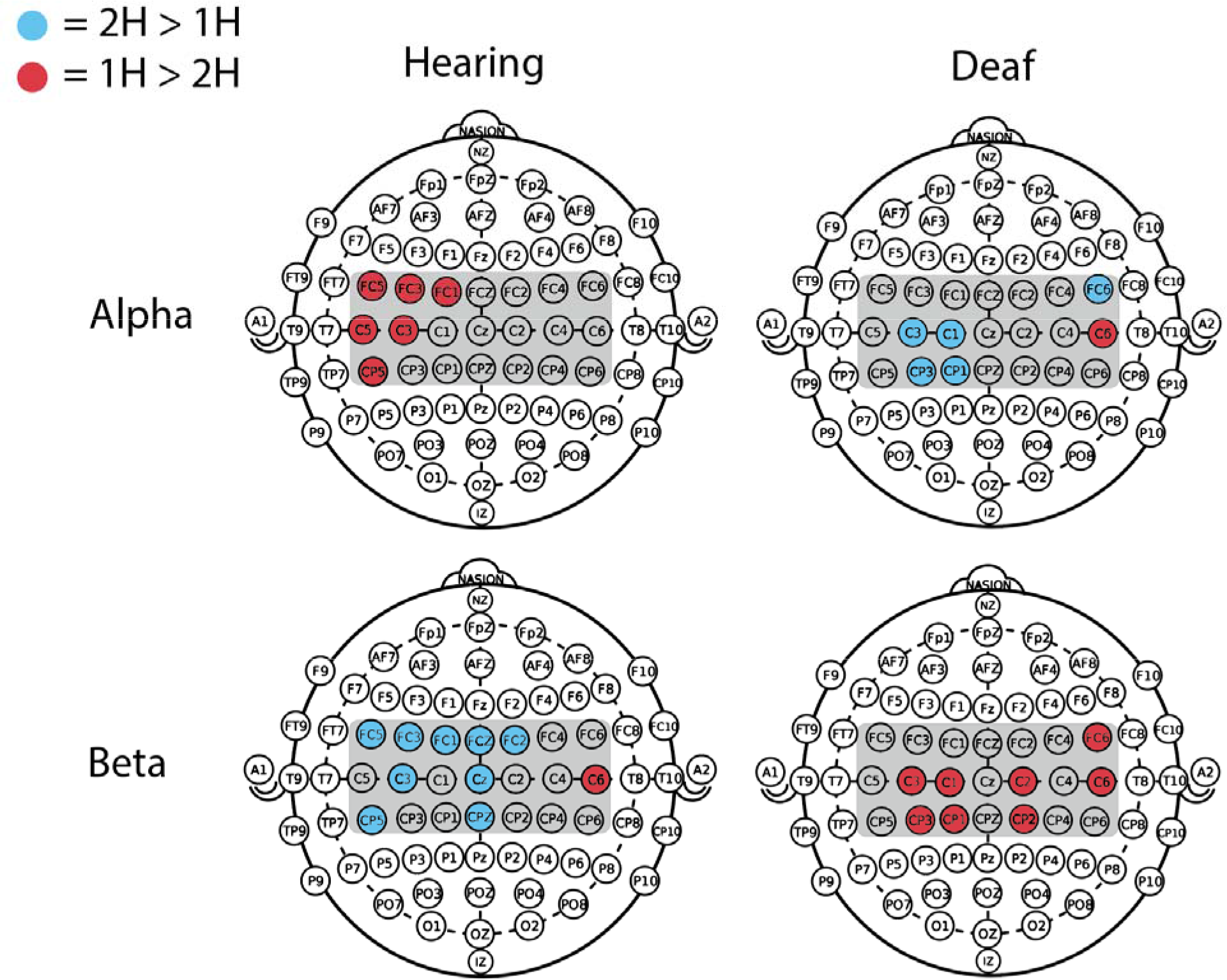
Electrode maps indicating significant differences in alpha (first row) and beta (second row) desynchronization for Hearing and Deaf groups. Gray shaded area indicates the central ROI. Blue shading indicates sites where observation of 2H signs led to more desynchronization than 1H signs. Red shading indicates sites where observation of 1H signs led to more desynchronization than 2H signs.

**Figure 3.**
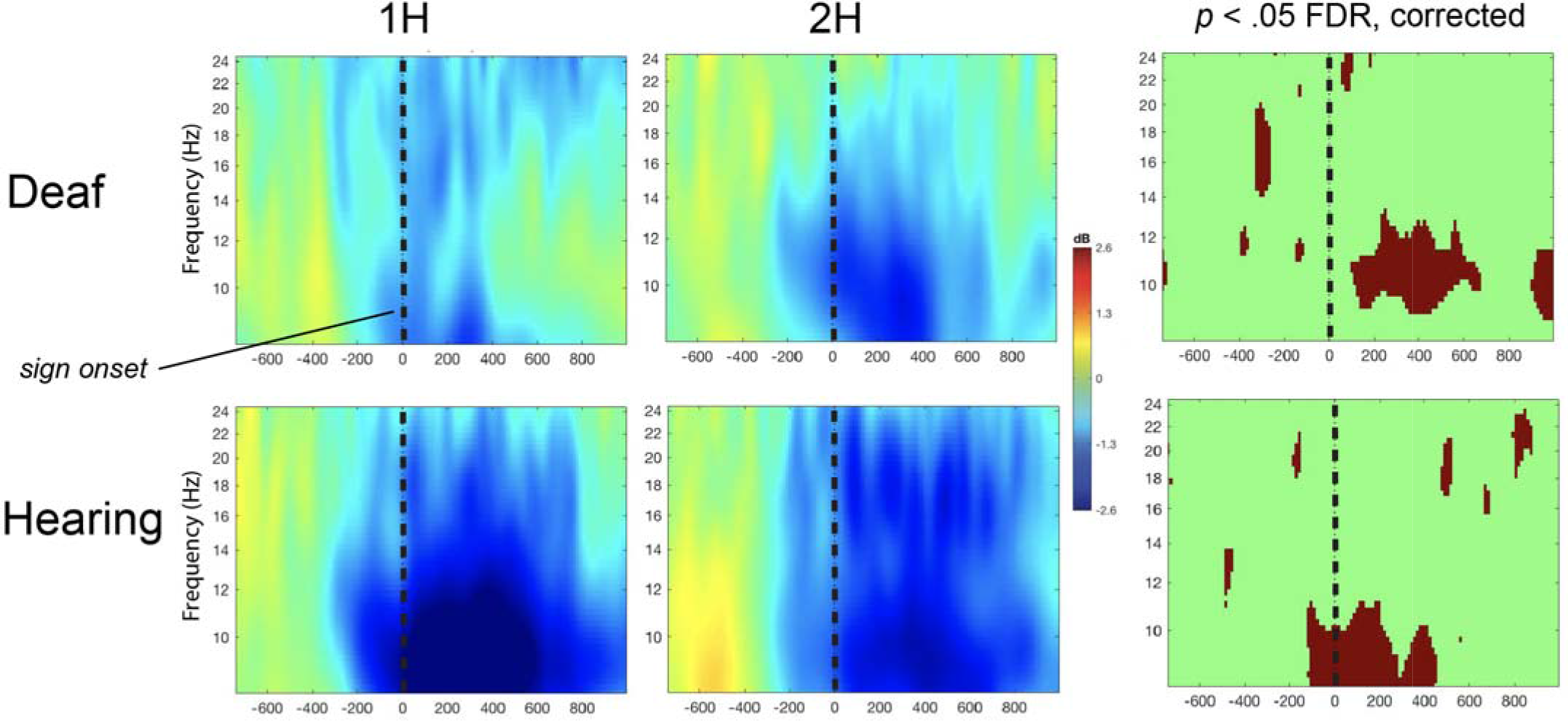
Alpha and beta power during presentation of 1H and 2H signs. Time-frequency plots of electrode C3 in response to observed signs. Cool colors indicate a decrease in power, warm colors indicate an increase in power. The first two columns show responses to 1H and 2H signs, respectively. The third column shows *p* values of the comparison of the first two panels, with the false discovery rate (FDR) correction applied.

In the Hearing group, six central electrodes showed significant effects of condition (1H v 2H) on alpha power (8-13 Hz) during sign perception; (*p* < .05, FDR corrected). Each of these electrodes showed greater alpha desynchronization in the 1H condition as compared to the 2H condition (see Figure 2). Ten electrodes showed significant effects of condition (1H v. 2H) on beta power (14-25 Hz) during sign perception (*p* < .05, FDR corrected). For nine electrodes, there was greater beta desynchronization in the 2H condition as compared to the 1H condition. One electrode showed the opposite effect within beta.

### 3.3. Occipital electrode analysis

In order to ascertain whether the observed results were due to general visual attention mechanisms we also conducted paired t-tests across the entire epoch (~-1000 ms to 1500 ms) comparing alpha and beta EEG responses to 1H and 2H stimuli at occipital electrodes (O1 and O2) overlying the visual cortex. The Deaf group showed no significant differences between 1H and 2H stimuli (*p* > .05, FDR corrected) at either occipital electrode. The Hearing group showed greater alpha desynchronization for 1H signs at both occipital electrode sites around −300 ms prior to sign onset. During this time, the signing model was present on the screen and was initiating the sign.

## 4. Discussion

The goal of the current study was to investigate the extent to which deaf signers engage sensorimotor cortices during perception of ASL signs, and to compare this activity to that of hearing non-signers. To that end, we designed a study asking how sensorimotor alpha and beta rhythms are influenced by observation of 1-handed and 2-handed signs in deaf signers and hearing non-signers. Two of the main findings of the present study are as follows: 1) Hearing non-signing participants showed greater sensorimotor cortex activity in response to signs compared to deaf signing participants; 2) Sensorimotor system activity in both deaf signers and hearing non-signers is sensitive to the gross sensorimotor characteristics of observed signs, suggesting that action mirroring processes are involved in sign perception for both groups.

### 4.1. Deaf signers vs. Hearing non-signers

Our first aim was to compare sensorimotor system activity in deaf signers and hearing non-signers as they perceived individual ASL signs. We predicted, based on findings in the action-expertise literature, that deaf signers would show greater alpha/beta rhythm desynchronization during ASL perception. However, the full scalp analyses showed that Hearing non-signers showed more alpha and beta desynchronization during sign observation, in contrast to our prediction. In fact, our data support the conclusion drawn by studies using PET and fMRI in that hearing non-signers recruit AON systems during passive ASL observation moreso than deaf fluent signers (Corina et al., 2007; Emmorey et al., 2010). This body of work suggests that the human AON is selective of action type, and therefore does not follow a linear model of experience-dependent activation for simply any action. Babiloni et al. (2009) examined the AON in response to non-athletes, amateur karate athletes, and elite karate athletes’ perception of karate action. When compared to non-athletes, elite karate athletes showed less AON recruitment, while when compared to non-athletes and elite athletes, amateur athletes had moderate AON recruitment. Similarly, Gardner et al. (2017) suggests a non-linear relationship between experience and AON involvement during subsequent observation. Our current results support this notion, demonstrating that at least with sign language experience, greater action experience does not lead to greater involvement of the sensorimotor cortex during observation.

The “neural efficiency” hypothesis states that with greater motor expertise in a given domain, there will be more efficient cortical functioning (Babiloni et al., 2009; Gardner et al., 2017; Haier, Siegel, MacLachlan, & Soderling, 1992; Rypma, Berger, & Desposito, 2002). It is possible that deaf fluent signers implement a top-down processing procedure to categorize perceived actions as either linguistic or non-linguistic. This top-down theory is supported by work in which deaf fluent signers recruit canonical language areas during sign language observation and middle-occipital/temporal regions (areas sensitive to human body form and movement) during observation of non-linguistic actions (Corina et al., 2007; Emmorey et al., 2010). Although previous work involving non-linguistic complex actions (i.e. dancing; Calvo-Merino et al., 2005; Orgs et al., 2008) has reported an increase in AON activity during subsequent observation for experts, our analyses on the effect of group support the existence of a “neural efficiency” type model for linguistic action experience and AON involvement. Thus, sign language processing in the AON does not have a linear experience-dependent relationship. ASL experience is unique because a familiarity with ASL involves not only motor knowledge, but also linguistic and conceptual knowledge. For a fluent signer, each sign carries a meaning, and sequences of signs include grammar, tone, and communicative intent. Thus, the results reported here likely reflect the accumulation of both the sensorimotor aspects of ASL action knowledge, and the linguistic conceptual knowledge that is embedded within. Conceptual and sensorimotor action knowledge are processed differently from one another in hearing populations (Gerson, Meyer, Hunnius, & Bekkering, 2017; Quandt & Chatterjee, 2015), and so the conceptual and motor effects of ASL experience upon subsequent perception are difficult to disentangle.

### 4.2. Sensorimotor Characteristics of Signs

Aside from the comparison between hearing non-signers and deaf signers, we wanted to assess whether deaf signers draw upon their own sensorimotor representations of observed signs to any extent. To test this, we varied the gross motor characteristics of the observed signs by comparing signs that use two hands to signs which are carried out on one hand only. We conducted our ROI analyses at electrodes overlying sensorimotor cortices in Deaf and Hearing groups. Both Hearing and Deaf groups showed differential responses to 1H and 2H signs, but the specifics of the oscillatory response profiles were quite complex. Our prediction of greater alpha and beta rhythm desynchronization for 2H signs in both groups was not supported. Rather, observation of 2H signs resulted in greater alpha desynchronization only for the Deaf group, and greater beta desynchronization only for the Hearing group. In contrast, observation of 1H signs resulted in greater alpha desynchronization for the Hearing group, and greater beta desynchronization in the Deaf group. The alpha and beta bands often show different effects in response to sensorimotor processing, and these differences may be attributable to different aspects of stimulus processing (Weiss & Mueller, 2012).

We conducted between-group analyses at occipital electrodes in order to test whether our observed effects were driven by visual attention. The Deaf group showed no significant differences between 1H and 2H within occipital electrode sites, and the Hearing group showed only a transient difference between conditions well before sign onset occurred. Thus, the effects we present here cannot be attributed to the occipital alpha rhythm.

### 4.3. Alpha-range effects

Producing a two-handed sign requires involvement of more of the primary sensorimotor cortices than the production of a one-handed sign (Emmorey et al., 2016). This activation is attributed to greater sensorimotor demands of using two limbs simultaneously as compared to one, as the same effects are seen with simple one- and two-handed movements (Toyokura et al., 2002). Due to this previous work, as well as work from the broader action mirroring field, we predicted that there would be significantly greater alpha desynchronization during observation of 2H signs than 1H signs. A cluster of electrodes for the Deaf group does support this prediction (see Figure 2), whereas in the Hearing group it was 1H signs which resulted in greater desynchronization. Taken together, these results suggest that the sensorimotor cortex is sensitive to the gross sensorimotor characteristics of observed actions, for both Deaf and Hearing groups. However, the flipped direction of the alpha rhythm desynchronization in response to the 1H/2H conditions across the two groups was not expected.

Alpha range activity over the sensorimotor cortices is broadly known to be sensitive to action mirroring processes (Aridan, Ossmy, Buaron, Reznik, & Mukamel, 2018; Arnstein, Cui, Keysers, Maurits, & Gazzola, 2011b; Bowman et al., 2017; Debnath et al., 2019). We expected the greater sensorimotor demands of a two-handed sign to result in greater sensorimotor cortex activity, and this is what we observed in the Deaf group, who are effectively action experts in the domain of ASL. However, Hearing non-signers showed greater alpha desynchronization over the left sensorimotor cortex for 1H signs. It is possible that their unfamiliarity with the observed stimuli led to this pattern of results. Hearing non-signers’ increased sensorimotor activation for 1H signs may be related to their increased ability to imitate simpler one-handed signs. For a Hearing non-signer, ASL signs are devoid of meaning, and one-handed signs may be simpler compared to signs produced using two hands. 1H signs typically use less signing space, use only one hand shape, and contain fewer locations than two handed signs, making them easier to process if there is no prior familiarity with the movement, as is the case with the hearing non-signers. Prior work shows that hearing non-signers show more mirror system activity when observing non-sign gestures than when observing sign language (MacSweeney et al., 2004). Thus, it is possible that greater alpha desynchronization we found for Hearing non-signers during observation of 1H signs is due to their increased ability to imitate them. Some work has shown that the more meaningless a hand movement is to the observer, the greater the alpha desynchronization (Streltsova, Berchio, Gallese, & Umilta, 2010), which again fits with our observed results. The effects of sign meaning and complexity upon perceptual processes is an area which would benefit from further research.

Deaf signers recruit sensorimotor cortices differently during execution of 1H and 2H signs (Emmorey et al., 2016). Given that greater alpha desynchronization in response to 2H signs happened primarily within the first 600 ms post-stimulus onset, it is possible that a type of inhibitory rebound is taking place. Desynchronization in alpha-range frequencies soon after stimulus onset, followed by a period of increased synchronization (i.e. ‘inhibitory rebound’) is said to reflect inhibition of MNS activity (Babiloni et al., 2002; Pfurtscheller, Brunner, Schlögl, & Lopes da Silva, 2006; Schuch, Bayliss, Klein, & Tipper, 2010). Thus, it is possible that the Deaf signing group exhibited motor resonance during 2H sign observation, closely followed by inhibitory control (i.e. mu synchronization) in order to regulate automatic imitation processes.

For both groups, the differences in sensorimotor cortex activity in response to sign characteristics were in primarily over the left sensorimotor cortex. We had expected that we would see mu rhythm sensitivity to the sensorimotor characteristics of signs over right hemishpere areas during 2H sign observation, as the primary difference between 1H and 2H signs is the involvement of the left hand. Our pattern of results may be explained by the fact that during action observation, the same bilateral responses are not seen, due to the lack of explicit motor planning involved in the task, as has been found in prior work (Avanzini et al., 2012; Quandt & Kubicek, 2018). Overall, the results seen in the alpha range EEG frequencies suggest that while both Deaf and Hearing groups were sensitive to the observed sensorimotor characteristics of signs, only the Deaf group showed the predicted effect of more sensorimotor cortex activity in response to signs which had greater sensorimotor demands.

### 4.4. Beta-range effects

Based on previous sign execution and action simulation research, we predicted there would be significantly greater beta desynchronization during the observation of 2H signs for both Deaf and Hearing groups due to the greater sensorimotor demands of performing an action with two hands. In fact, we found that while the Hearing group showed more beta desynchronization in response to 2H signs, the opposite pattern was observed for the Deaf group. Thus, we found our prediction to only be true for the Hearing group. While many researchers show greater beta desynchronization in response to greater sensorimotor engagement, there is a growing body of evidence demonstrating the opposite effect: beta desynchronization in response to smaller or less demanding movements (Cheyne et al., 2003; Neuper & Pfurtscheller, 2001; Pfurtscheller, Krausz, & Neuper, 2001; Quandt et al., 2013; Spitzer & Haegens, 2017). A recent study from our laboratory showed increased beta desynchronization when deaf signers read English translations of one-handed signs compared to 2H signs (Quandt & Kubicek, 2018). The current results for the Deaf group are in line with this prior finding. In both studies, the greater sensorimotor cortex activity in response to 1H stimuli is likely due to a combination of vicarious motor processing and inhibition of automatic imitation. Our current results contribute evidence to the growing understanding that activity within the beta rhythm reflects complex interactions between language and sensorimotor processing (Spitzer & Haegens, 2017; Weiss & Mueller, 2012).

Overall, our current results suggest that varying gross sensorimotor characteristics of observed signs differentially engage observers’ sensorimotor brain regions. In contrast to prior work demonstrating little or no involvement of action mirroring processes during signers’ observation of sign language, we demonstrate that deaf signers do draw upon their own sensorimotor cortices during sign observation, in a manner which is sensitive to the physical and kinematic properties of the signs they are seeing. It may be that the increased temporal resolution of EEG time-frequency analyses has allowed for a view of neural mechanisms which would be missed by hemodynamic neuroimaging methods. This may explain why our results reveal action mirroring processes in signers contrary to prior neuroimaging studies (Corina et al., 2007; Emmorey et al., 2010). However, given that the results presented here are the first time-frequency analyses of deaf signers viewing signs, we recognize the need for future work in this area to further differentiate the contributions of linguistic processing from those of action mirroring.

## 5. Future directions and conclusions

While all the deaf signers included in the current study were self-identified as fluent and the instructions for the procedure were given in American Sign Language, we did not statistically control for, or otherwise analyze, the age of ASL acquisition or educational background (i.e. mainstream, deaf school). Also, no group comprised of deaf non-signers or hearing signers was included in this design. Including a deaf non-signing group would control for effects of hearing loss, while hearing signers would control for effects of sign language knowledge. Disentangling effects of hearing status and language experience would shed further light on role the AON plays in amodal language facilitation. Given the complex and likely non-linear nature of the relationship between action experience and action mirroring, it is natural to wonder how varying amounts of sign language experience would affect subsequent observation. Future work should examine the profile of new ASL signers, or attempt to give novices controlled training experiences with ASL. Only through work such as this will we be able to paint a fuller picture of how ASL knowledge changes the neural substrates of action perception.

The goal of the current work was to shed further light on the relationship between the Action Observation Network (AON) and sign language perception. By framing signs as complex actions and controlling for gross motor differences within the action, we designed this study to be similar to other action observation research (Calvo-Merino et al., 2005; Cross, Hamilton, & Grafton, 2006b; Gardner et al., 2015; Orgs et al., 2008). We aimed to answer two questions: 1) How do Deaf and Hearing groups differ in sensorimotor EEG rhythm activity during perception of ASL signs; and 2) are both groups sensitive to the gross sensorimotor characteristics of observed signs? The second of these questions allowed us to explore the possibility that even if non-signers show more AON recruitment during sign observation, the signers may still be drawing upon sensorimotor cortices when perceiving signs.

Full scalp map analyses across Deaf and Hearing groups yield results in line with previous sign language/AON research (Corina et al., 2007; Emmorey et al., 2010); hearing non-signers show greater sensorimotor EEG desynchronization than deaf signers during the observation of sign language. Less sensorimotor engagement for the Deaf group is attributed to top-down processing involving the Deaf group transferring the cognitive load to more canonical language areas upon perception, as opposed to processing in typical action observation areas. Work investigating the interaction between the AON and sign language has strongly suggested a lack of MNS recruitment for deaf signers during sign language observation. Time-frequency analyses over the central ROI demonstrate for the first time that the mirroring properties of the AON are recruited when deaf signers observe signs, as demonstrated through the differential mu rhythm desynchronization for 1H and 2H signs. By including conditions that facilitate the comparison of sign types we demonstrate that future sign language/AON research would benefit from poising signs as complex actions, resulting in a better understanding of overall AON functioning. Significant work still needs to be done to explicate the relationship between these complex linguistic actions and the AON, including further delineation of sign type and varying groups of sign language users (i.e. deaf, hearing). Analyses following this study will illuminate the boundaries of the AON in human language, informing on MNS roles within human social cognition.

## Acknowledgements

The authors are grateful to Naseem Majrud, Taylor Wardle, and Athena Willis for assistance in data collection. We are also thankful for each of the individuals who took the time to participate in this research study. We acknowledge the support of Gallaudet University, NSF/Gallaudet University’s Science of Learning Center, VL2 (Visual Language and Visual Learning), and NSF Grant #1839379 to Lorna Quandt.

## References

Aridan, N., Ossmy, O., Buaron, B., Reznik, D., & Mukamel, R. (2018). Suppression of EEG mu rhythm during action observation corresponds with subsequent changes in behavior. Brain Research, 1691, 55–63. http://doi.org/10.1016/j.brainres.2018.04.013

Arnstein, D., Cui, F., Keysers, C., Maurits, N. M., & Gazzola, V. (2011a). Mu Suppression during Action Observation and Execution Correlates with BOLD in Dorsal Premotor, Inferior Parietal, and SI Cortices. Journal of Neuroscience, 31(40), 14243–14249. http://doi.org/10.1523/jneurosci.0963-11.2011

Arnstein, D., Cui, F., Keysers, C., Maurits, N. M., & Gazzola, V. (2011b). Mu-Suppression during Action Observation and Execution Correlates with BOLD in Dorsal Premotor, Inferior Parietal, and SI Cortices. Journal of Neuroscience, 31(40), 14243–14249. http://doi.org/10.1523/JNEUROSCI.0963-11.2011

Avanzini, P., Fabbri-Destro, M., Dalla Volta, R., Daprati, E., Rizzolatti, G., & Cantalupo, G. (2012). The Dynamics of Sensorimotor Cortical Oscillations during the Observation of Hand Movements: An EEG Study. PLoS ONE, 7(5), e37534–10. http://doi.org/10.1371/journal.pone.0037534

Babiloni, C., Babiloni, F., Carducci, F., Cincotti, F., Cocozza, G., Del Percio, C., et al. (2002). Human Cortical Electroencephalography (EEG) Rhythms during the Observation of Simple Aimless Movements: A High-Resolution EEG Study. NeuroImage, 17(2), 559–572. http://doi.org/10.1006/nimg.2002.1192

Babiloni, C., Del Percio, C., Rossini, P. M., Marzano, N., Iacoboni, M., Infarinato, F., et al. (2009). Judgment of actions in experts: A high-resolution EEG study in elite athletes. NeuroImage, 45(2), 512–521. http://doi.org/10.1016/j.neuroimage.2008.11.035

Babiloni, C., Marzano, N., Infarinato, F., Iacoboni, M., Rizza, G., Aschieri, P., et al. (2010). “Neural efficiency” of experts’ brain during judgment of actions: A high-resolution EEG study in elite and amateur karate athletes. Behavioural Brain Research, 207(2), 466–475. http://doi.org/10.1016/j.bbr.2009.10.034

Benjamini, Y., Drai, D., Elmer, G., & Kafkafi, N. (1995). Controlling the false discovery rate in behavior genetics research. Journal of the Royal Statistical Society.

Bowman, L. C., Bakermans-Kranenburg, M. J., Yoo, K. H., Cannon, E. N., Vanderwert, R. E., Ferrari, P. F., et al. (2017). The mu-rhythm can mirror: Insights from experimental design, and looking past the controversy, 96, 121–125. http://doi.org/10.1016/j.cortex.2017.03.025

Braadbaart, L., Williams, J. H. G., & Waiter, G. D. (2013). Do mirror neuron areas mediate mu rhythm suppression during imitation and action observation? International Journal of Psychophysiology, 89(1), 1–7. http://doi.org/10.1016/j.ijpsycho.2013.05.019

Calvo-Merino, B., Glaser, D. E., Grèzes, J., Passingham, R. E., & Haggard, P. (2005). Action Observation and Acquired Motor Skills: An fMRI Study with Expert Dancers. Cerebral Cortex, 15(8), 1243–1249. http://doi.org/10.1093/cercor/bhi007

Caselli, N. K., Sehyr, Z. S., Cohen-Goldberg, A. M., & Emmorey, K. (2016). ASL-LEX: A lexical database of American Sign Language. http://doi.org/10.3758/s13428-016-0742-0

Caspers, S., Zilles, K., Laird, A. R., & Eickhoff, S. B. (2010). ALE meta-analysis of action observation and imitation in the human brain. NeuroImage, 50(3), 1148–1167. http://doi.org/10.1016/j.neuroimage.2009.12.112

Cheyne, D., Gaetz, W., Garnero, L., Lachaux, J.-P., Ducorps, A., Schwartz, D., & Varela, F. J. (2003). Neuromagnetic imaging of cortical oscillations accompanying tactile stimulation. Cognitive Brain Research, 17(3), 599–611. http://doi.org/10.1016/S0926-6410(03)00173-3

Corina, D., Chiu, Y.-S., Knapp, H., Greenwald, R., San Jose-Robertson, L., & Braun, A. (2007). Neural correlates of human action observation in hearing and deaf subjects. Brain Research, 1152, 111–129. http://doi.org/10.1016/j.brainres.2007.03.054

Cross, E. S., Cohen, N. R., Hamilton, A. F. de C., Ramsey, R., Wolford, G., & Grafton, S. T. (2012). Physical experience leads to enhanced object perception in parietal cortex: Insights from knot tying. Neuropsychologia, 50(14), 3207–3217. http://doi.org/10.1016/j.neuropsychologia.2012.09.028

Cross, E. S., Hamilton, A. F. de C., Kraemer, D. J. M., Kelley, W. M., & Grafton, S. T. (2009). Dissociable substrates for body motion and physical experience in the human action observation network. European Journal of Neuroscience, 30(7), 1383–1392. http://doi.org/10.1111/j.1460-9568.2009.06941.x

Cross, E. S., Hamilton, A. F. D. C., & Grafton, S. T. (2006). Building a motor simulation de novo: observation of dance by dancers. Neuroimage, 31(3), 1257–1267.

Cross, E. S., Liepelt, R., de C Hamilton, A. F., Parkinson, J., Ramsey, R., Stadler, W., & Prinz, W. (2011). Robotic movement preferentially engages the action observation network. Human Brain Mapping, 33(9), 2238–2254. http://doi.org/10.1002/hbm.21361

Debnath, R., Salo, V. C., Buzzell, G. A., Yoo, K. H., & Fox, N. A. (2019). Mu rhythm desynchronization is specific to action execution and observation: Evidence from time-frequency and connectivity analysis. NeuroImage, 184, 496–507. http://doi.org/10.1016/j.neuroimage.2018.09.053

Delorme, A., & Makeig, S. (2004). EEGLAB: an open source toolbox for analysis of single-trial EEG dynamics including independent component analysis. Journal of Neuroscience Methods.

Drew, A. R., Quandt, L. C., & Marshall, P. J. (2015). Visual influences on sensorimotor EEG responses during observation of hand actions. Brain Research, 1597(C), 119–128. http://doi.org/10.1016/j.brainres.2014.11.048

Emmorey, K., Mehta, S., McCullough, S., & Grabowski, T. J. (2016). The neural circuits recruited for the production of signs and fingerspelled words. Brain and language, 160, 30–41.

Emmorey, K., Xu, J., Gannon, P., Goldin-Meadow, S., & Braun, A. (2010). CNS activation and regional connectivity during pantomime observation: No engagement of the mirror neuron system for deaf signers. NeuroImage, 49(1), 994–1005. http://doi.org/10.1016/j.neuroimage.2009.08.001

Errante, A., & Fogassi, L. (2019). Parieto-frontal mechanisms underlying observation of complex hand-object manipulation. Scientific reports, 9(1), 348.

Fadiga, L., Fogassi, L., Pavesi, G., & Rizzolatti, G. (1995). Motor facilitation during action observation: a magnetic stimulation study. Physiology.org.

Fox, N. A., Bakermans-Kranenburg, M. J., Yoo, K., Bowman, L., Cannnon, E., Vanderwert, E., et al. (2016). Assessing human mirror activity with EEG mu rhythm: A meta-analysis. Psycnet.Apa.org. http://doi.org/10.1037/bul0000031.supp

Gardner, T., Aglinskas, A., & Cross, E. S. (2017). Using guitar learning to probe the Action Observation Network’s response to visuomotor familiarity. NeuroImage, 156, 174–189. http://doi.org/10.1016/j.neuroimage.2017.04.060

Gardner, T., Goulden, N., & Cross, E. S. (2015). Dynamic Modulation of the Action Observation Network by Movement Familiarity. Journal of Neuroscience, 35(4), 1561–1572. http://doi.org/10.1523/JNEUROSCI.2942-14.2015

Gazzola, V., Rizzolatti, G., Wicker, B., & Keysers, C. (2007). The anthropomorphic brain: The mirror neuron system responds to human and robotic actions. NeuroImage, 35(4), 1674–1684. http://doi.org/10.1016/j.neuroimage.2007.02.003

Gerson, S. A., Meyer, M., Hunnius, S., & Bekkering, H. (2017). Unravelling the contributions of motor experience and conceptual knowledge in action perception: A training study. Scientific Reports, 1–10. http://doi.org/10.1038/srep46761

Grèzes, J., Armony, J. L., Rowe, J., & Passingham, R. E. (2003). Activations related to “mirror” and ‘canonical’ neurones in the human brain: an fMRI study. NeuroImage, 18(4), 928–937. http://doi.org/10.1016/S1053-8119(03)00042-9

Haier, R. J., Siegel, B. V., Jr, MacLachlan, A., & Soderling, E. (1992). Regional glucose metabolic changes after learning a complex visuospatial/motor task: a positron emission tomographic study. Brain Research.

Hardwick, R. M., Caspers, S., Eickhoff, S. B., & Swinnen, S. P. (2017). Neural Correlates of Motor Imagery, Action Observation, and Movement Execution: A Comparison Across Quantitative Meta-Analyses, 1–50. http://doi.org/10.1101/198432

Hari, R. (2006). Action–perception connection and the cortical mu rhythm. In Event-Related Dynamics of Brain Oscillations (Vol. 159, pp. 253–260). Elsevier. http://doi.org/10.1016/S0079-6123(06)59017-X

Hari, R., Forss, N., Avikainen, Kirveskari, Salenius, & Rizzolatti, G. (1998). Activation of human primary motor cortex during action observation: a neuromagnetic study. National Acad Sciences.

Hobson, H. M., & Bishop, D. V. M. (2017). The interpretation of mu suppression as an index of mirror neuron activity: past, present and future. Royal Society Open Science, 4(3), 160662–22. http://doi.org/10.1098/rsos.160662

Iacoboni, M. (2009). Imitation, Empathy, and Mirror Neurons. Annual Review of Psychology, 60(1), 653–670. http://doi.org/10.1146/annurev.psych.60.110707.163604

Liew, S.-L. (2013). Both novelty and expertise increase action observation network activity, 1–15. http://doi.org/10.3389/fnhum.2013.00541/abstract

MacSweeney, M., Campbell, R., Woll, B., Giampietro, V., David, A. S., McGuire, P. K., et al. (2004). Dissociating linguistic and nonlinguistic gestural communication in the brain. NeuroImage, 22(4), 1605–1618. http://doi.org/10.1016/j.neuroimage.2004.03.015

MacSweeney, M., Woll, B., & Campbell, R. (2002). Neural correlates of British sign language comprehension: spatial processing demands of topographic language. Journal of Cognitive Neuroscience, 14(7), 1064–1075. http://doi.org/10.1162/089892902320474517

Montgomery, K. J., Isenberg, N., & Haxby, J. V. (2007). Communicative hand gestures and object-directed hand movements activated the mirror neuron system. Social Cognitive and Affective Neuroscience, 2(2), 114–122. http://doi.org/10.1093/scan/nsm004

Muise-Hennessey, Tremblay, White, McWhinney, Zaini, Maessen, et al. (2016). Age of Onset and Duration of Deafness Drive Brain Organization for Biological Motion Perception in Non-Signers (pp. 1–58).

Muthukumaraswamy, S. D., Johnson, B. W., & McNair, N. A. (2004). Mu rhythm modulation during observation of an object-directed grasp. Cognitive Brain Research, 19(2), 195–201. http://doi.org/10.1016/j.cogbrainres.2003.12.001

Neuper, C., & Pfurtscheller, G. (2001). Evidence for distinct beta resonance frequencies in human EEG related to specific sensorimotor cortical areas. Clinical Neurophysiology.

Newman, A. J., Supalla, T., Fernandez, N., Newport, E. L., & Bavelier, D. (2015). Neural systems supporting linguistic structure, linguistic experience, and symbolic communication in sign language and gesture. Proceedings of the National Academy of Sciences, 112(37), 11684–11689. http://doi.org/10.1073/pnas.1510527112

Orgs, G., Dombrowski, J.-H., Heil, M., & Jansen-Osmann, P. (2008). Expertise in dance modulates alphabeta event-related desynchronization during action observation. European Journal of Neuroscience, 27(12), 3380–3384. http://doi.org/10.1111/j.1460-9568.2008.06271.x

Perry, A., & Bentin, S. (2009). Mirror activity in the human brain while observing hand movements: A comparison between EEG desynchronization in the μ-range and previous fMRI results. Brain Research, 1–7. http://doi.org/10.1016/j.brainres.2009.05.059

Pfurtscheller, G., Brunner, C., Schlögl, A., & Lopes da Silva, F. H. (2006). Mu rhythm (de)synchronization and EEG single-trial classification of different motor imagery tasks. NeuroImage, 31(1), 153–159. http://doi.org/10.1016/j.neuroimage.2005.12.003

Pfurtscheller, G., Krausz, G., & Neuper, C. (2001). Mechanical stimulation of the fingertip can induce bursts of β oscillations in sensorimotor areas. Clinical Neurophysiology.

Pineda, J. A., Allison, B. Z., & Vankov, A. (2000). The effects of self-movement, observation, and imagination on μ rhythms and readiness potentials (RP’s): toward a brain-computer interface (BCI). IEEE Transactions on Rehabilitation Engineering, 8(2), 219–222. http://doi.org/10.1109/86.847822

Quandt, L. C., & Chatterjee, A. (2015). Rethinking actions: implementation and association. Wiley Interdisciplinary Reviews: Cognitive Science, 6(6), 483–490. http://doi.org/10.1002/wcs.1367

Quandt, L. C., & Kubicek, E. (2018). Sensorimotor characteristics of sign translations modulate EEG when deaf signers read English. Brain and Language, 187, 9–17. http://doi.org/10.1016/j.bandl.2018.10.001

Quandt, L. C., & Marshall, P. J. (2014). The effect of action experience on sensorimotor EEG rhythms during action observation. Neuropsychologia, 56, 401–408. http://doi.org/10.1016/j.neuropsychologia.2014.02.015

Quandt, L. C., Marshall, P. J., Bouquet, C. A., & Shipley, T. F. (2013). Somatosensory experiences with action modulate alpha and beta power during subsequent action observation. Brain Research, 1534, 55–65. http://doi.org/10.1016/j.brainres.2013.08.043

Quandt, L. C., Marshall, P. J., Shipley, T. F., Beilock, S. L., & Goldin-Meadow, S. (2012). Sensitivity of alpha and beta oscillations to sensorimotor characteristics of action: An EEG study of action production and gesture observation. Neuropsychologia, 50(12), 2745–2751. http://doi.org/10.1016/j.neuropsychologia.2012.08.005

Ritter, P., Moosmann, M., & Villringer, A. (2009). Rolandic alpha and beta EEG rhythms’ strengths are inversely related to fMRI-BOLD signal in primary somatosensory and motor cortex. Human Brain Mapping, 30(4), 1168–1187. http://doi.org/10.1002/hbm.20585

Rizzolatti, G., & Fogassi, L. (2014). The mirror mechanism: recent findings and perspectives. Philisophical Transactions of the Royal Society. http://doi.org/10.1098/rstb.2013.0420

Rizzolatti, G., & Sinigaglia, C. (2010). The functional role of the parieto-frontal mirror circuit: interpretations and misinterpretations, 1–11. http://doi.org/10.1038/nrn2805

Rizzolatti, G., Fogassi, L., & Gallese, V. (2001). Neurophysiological mechanisms underlying the understanding and imitation of action. Nature.com. http://doi.org/10.1038/35090060

Rypma, B., Berger, J., & Desposito, M. (2002). The Influence of Working-Memory Demand and Subject Performance on Prefrontal Cortical Activity, 1–11.

Schuch, S., Bayliss, A. P., Klein, C., & Tipper, S. P. (2010). Attention modulates motor system activation during action observation: evidence for inhibitory rebound. Experimental Brain Research, 205(2), 235–249. http://doi.org/10.1007/s00221-010-2358-4

Spitzer, B., & Haegens, S. (2017). Beyond the Status Quo: A Role for Beta Oscillations in Endogenous Content (Re)Activation. Eneuro, 4(4), ENEURO.0170–17.2017–49. http://doi.org/10.1523/ENEURO.0170-17.2017

Streltsova, A., Berchio, C., Gallese, V., & Umilta, M. A. (2010). Time course and specificity of sensory-motor alpha modulation during the observation of hand motor acts and gestures: a high density EEG study. Experimental Brain Research, 205(3), 363–373. http://doi.org/10.1007/s00221-010-2371-7

Toyokura, M., Muro, I., Komiya, T., Obara. (2002). Activation of Pre–Supplementary Motor Area (SMA) and SMA Proper During Unimanual and Bimanual Complex Sequences: An Analysis Using Functional Magnetic …. Journal of Neuroimaging.

Ulloa, E., & Pineda, J. (2007). Recognition of point-light biological motion: Mu rhythms and mirror neuron activity, 183(2), 188–194. http://doi.org/10.1016/j.bbr.2007.06.007

van der Gaag, C., Minderaa, R. B., & Keysers, C. (2007). Facial expressions: What the mirror neuron system can and cannot tell us. Social Neuroscience, 2(3-4), 179–222. http://doi.org/10.1080/17470910701376878

Weiss, S., & Mueller, H. (2012). “Too many betas do not spoil the broth”: the role of beta brain oscillations in language processing. Frontiers in Psychology, 1–15. http://doi.org/10.3389/fpsyg.2012.00201/abstract

